# Targeting the ferroptosis pathway: A novel compound, AZD1390, protects the brain after ischemic stroke

**DOI:** 10.1101/2025.02.22.639635

**Authors:** Han Kyu Lee, Chao-Chieh Lin, Denise E. Dunn, Yubin Chen, Ssu-Yu Chen, Douglas A. Marchuk, Scott R. Floyd, Jen-Tsan Chi

## Abstract

**Background:** Ferroptosis is an iron-dependent form of regulated cell death driven by lipid peroxidation. This process has been implicated in various diseases, including ischemic stroke. Ischemic stroke leads to oxidative stress, iron overload, and reactive oxygen species (ROS) accumulation, which collectively may trigger ferroptotic neuronal cell death. However, the regulatory mechanisms of ferroptosis in stroke remain poorly understood. Previous studies have identified ataxia telangiectasia mutated (ATM), a DNA damage kinase, as a critical regulator of ferroptosis. However, the therapeutic potential of this discovery remains unknown.

**Methods:** We investigated the effect of ATM inhibitors, including the brain-penetrant AZD1390, on ferroptosis using *in vitro, ex vivo*, and *in vivo* models of ischemic stroke. Our analysis included assessments of cell viability, lipid peroxidation, ferroptosis marker expression, and infarct volume.

**Result:** ATM inhibitors significantly alleviated ferroptosis-induced cell death in cultured cells and *ex vivo* murine brain slice cultures. In the oxygen-glucose deprivation (OGD) stroke model, treatment with AZD1390 reduced the expression of ferroptosis markers (xCT and PTGS2) and diminished neuronal cell death in rat and mouse brain slices. Furthermore, in a mouse model of ischemic stroke, AZD1390 decreased infarct volume confirming its therapeutic efficacy *in vivo*.

**Conclusions:** This study identifies ferroptosis as a critical mechanism in ischemic stroke-induced neuronal cell death and highlights ATM inhibition, particularly with AZD1390, as a promising therapeutic candidate for mitigating stroke-associated damage. Targeting ferroptosis may provide a translationally relevant strategy to mitigate neuronal injury and improve clinical outcomes for stroke patients.

## Introduction

Ferroptosis, a regulated form of cell death driven by iron-dependent lipid peroxidation, has emerged as a critical mechanism implicated in various diseases ^1,2^. Initially identified as a mode of cell death in RAS-mutated cancer cells, ferroptosis is characterized by iron dependency, lipid peroxidation, and mitochondrial shrinkage ^1^. Its regulation involves key players such as glutathione peroxidase 4 (GPX4) and ferroptosis suppressor protein 1 (FSP1), which mitigate lipid peroxidation and protect against cell death ^3,4^. Dysregulated ferroptosis has been linked to diverse pathological conditions, including tumors, acute kidney injury, neurological disorders, and ischemia/reperfusion injury. Importantly, recent studies have identified ferroptosis as a pivotal neuronal cell death pathway triggered by ischemic injury, highlighting its relevance in ischemic stroke ^5,6^.

Ischemic stroke, caused by disrupted blood flow to the brain, results in oxidative stress, iron overload, and the accumulation of reactive oxygen species (ROS), all of which synergistically induce ferroptotic cell death ^5,6^. This process exacerbates neuronal damage and worsens stroke outcomes, highlighting ferroptosis as a potential therapeutic target. Despite its significance, the underlying regulatory mechanisms of ferroptosis in stroke remain incompletely understood.

Although inhibiting ferroptosis presents a promising therapeutic approach for ischemic stroke, the current ferroptosis inhibitors are not yet suitable for clinical application. To uncover key regulators of ferroptosis, we previously conducted forward kinome screens for modifiers of ferroptotic responses in cells, highlighting ataxia telangiectasia mutated (ATM) kinase as a top candidate ^7^. Our findings demonstrate that ATM inhibition reduces labile iron levels by inducing the expression of iron-sequestering proteins, such as ferritin and ferroportin, mediated through metal regulatory transcription factor 1 (MTF1) ^7^. This result has been further validated by other independent studies ^8,9^.

Given the availability of numerous ATM inhibitors developed for preclinical and clinical studies, we propose that ATM inhibition represents a promising translational approach to mitigating ferroptosis in ischemic stroke. For example, AZD1390, an orally bioavailable and brain-penetrant ATM inhibitor, has demonstrated efficacy in glioma ^10^ and is currently being evaluated in clinical trials (NCT03423628). In this study, we further investigated the therapeutic potential of ATM inhibition in regulating ferroptosis in the context of ischemic stroke. Our findings reveal that inhibiting ATM with brain-penetrating inhibitor AZD1390 significantly alleviates brain infarction in a model of ischemic stroke. These results underscore the therapeutic potential of targeting ferroptosis pathways as a physiologically relevant and promising translation pathway for ischemic stroke in humans.

## Materials and Methods

**Animals**. All inbred mouse strains were obtained from the Jackson Laboratory (Bar Harbor, ME), and then bred locally to generate mice used in all experiments. Sprague-Dawley (SD) rats for *ex vivo* brain slice culture experiments were obtained from Charles River (Wilmington, MA). 12 ± 1 week-old mice (both male and female animals) were used for pMCAO and postnatal day 7 mice and rats were used for *ex vivo* brain slice culture. All animal study procedures were conducted under protocols approved by the Duke University IACUC in accordance with NIH guidelines.

### AZD1390 preparation

Pharmaceutical-grade AZD1390 for *in vivo* studies was obtained directly from AstraZeneca and was prepared as recommended by the manufacture’s formulation and doses for *in vivo* mouse studies. Briefly, a dose of 20 mg/kg was suspended in a vehicle consisting of 0.5% hydroxypropyl methylcellulose (HPMC) and 0.1% Tween-80 in water.

### Lipid Peroxidation Assay

Lipid peroxidation was evaluated using the C11-BODIPY dye (ThermoFisher, D3861). Cells were treated with either vehicle or erastin with/without various doses of AZD1390 for 16 hours. Lipid peroxidation levels were analyzed using flow cytometry (FACSCanto II, BD Biosciences).

### Preparation of brain slices and oxygen-glucose deprivation (OGD)

Preparation of cortical brain slice explants was performed as previously described ^11,12^. Coronal brain slices were prepared from P7 rats and mice. Under sterile conditions, brains were dissected and cut into 250 μM hemi-coronal slices on a vibratome in a chilled culture medium. For OGD, slices were suspended at 34°C for 4.5 minutes of glucose-free N_2_-bubbled artificial CSF. Brain slices were plated into 12-well plates in interface configuration atop a solid culture medium made by adding 0.5% agarose. After explanting the brain slices, plates were placed for recovery at 30°C for 30 minutes in a humidified incubator under under 5% CO_2_. Gold particles (1.6 μM) coated with plasmids expressing YFP were introduced into the brain slices by biolistic transfection using a Helios Gene Gun (Bio-Rad). Slice cultures were maintained at 37 °C for 24 hours in a humidified incubator under 5% CO_2_.

### Permanent middle cerebral artery occlusion (pMCAO) and AZD1390 treatment

Focal cerebral ischemia was induced by direct permanent occlusion of the distal MCA as previously described ^13,14^. Two hours after pMCAO, animals received a 20 mg/kg dose of AZD1390 via oral gavage using a flexible feeding tube (Instech, Plymouth Meeting, PA). Mice were then returned to their cages and allowed free access to food and water in an air-ventilated room maintained at an ambient temperature of 25°C.

### Infarct volume measurement

Cerebral infarct volumes were measured 72 hours after distal permanent MCA occlusion as previously described ^13,14^.

### Statistical analyses

Statistical analyses were performed with GraphPad Prism (GraphPad Software, La Jolla, CA). Significant differences between datasets were evaluated using a two-tailed Student’s *t*-test comparisons between two groups or one-way ANOVA followed by Tukey’s multiple comparison test for more than two groups. Data are represented as the mean ± SEM. *p* < 0.05 was considered statistically significant. Detailed statistical analyses for all figures are provided in Table S1. Experimental details are included in the Supplemental Materials and Methods.

## Results

### ATM inhibitors protect against erastin-induced ferroptosis *in vitro*

To validate our previous finding that inhibiting ferroptosis enhances cell viability ^7^, we tested canonical ferroptosis inhibitors, deferoxamine (DFOA) and liproxstatin-1 (LPT1), and ATM inhibitors, KU55933 and KU60019. As expected, both ferroptosis inhibitors and ATM inhibitors significantly increased cell viability following erastin-induced ferroptosis (Fig 1A). These results confirm our previous observation that ATM plays a critical role in the ferroptosis pathway ^7^. We further investigated this by testing another ATM inhibitor, AZD1390, initially developed for cancer treatment by suppressing DNA double-strand break repair ^15^. AZD1390 is an orally bioavailable and brain-penetrant ATM inhibitor used for treating glioblastoma ^10^ and in a recent clinical trial (NCT03423628). We found AZD1390 dose-dependently reduced erastin-induced cell death (Fig 1B). We also measured membrane rupture and cytotoxicity to assess its effects on membrane integrity, observing a dose-dependent decrease in cell membrane rupture with AZD1390 treatment (Fig 1C). Additionally, erastin-induced lipid peroxidation, a hallmark of ferroptosis, was significantly reduced by AZD1390 treatment as measured by C11-BODIPY and flow cytometry (Fig 1D and E). These findings strongly support our hypothesis that inhibiting ATM prevents cell death by inhibiting ferroptosis.

**Figure 1.**
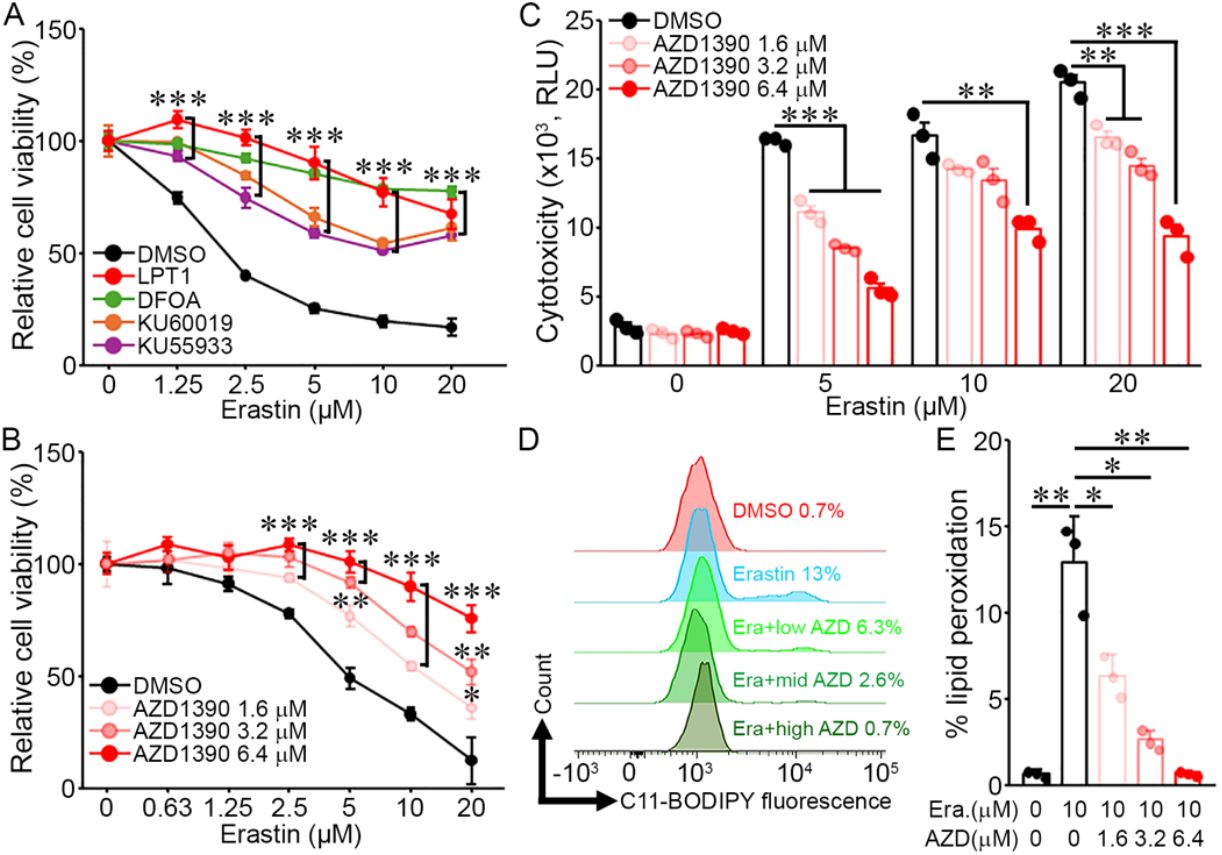
AZD1390 protects against erastin-induced ferroptosis in MDA-MB-231 cells. **(A)** The line graph shows cell viability of MDA-MB-231 cells. MDA-MB-231 cells were treated with increasing doses of erastin, either alone or in combination with deferoxamine (DFOA, 80μM), liproxstatin-1 (LPT1, 2μM), KU55933 (5μM), and KU60019 (1μM). The cell viability was quantified by CellTiter-Glo assay. **(B and C)** AZD1390 reduced erastin-induced ferroptosis in MDA-MB-231 cells in a dose-dependent manner. The cell viability and cytotoxicity were determined by CellTiter-Glo assay (B) and CellTox Green assay (C), respectively. **(D and E)** Erastin-induced lipid peroxidation (10μM, 18 hours) in MDA-MB-231 cells was reduced by AZD1390. Lipid peroxidation was assessed by C11-BODIPY staining (D) and quantified as the percentage of lipid peroxidation-positive cells (E). Data represent the mean ± SEM and statistical significance was determined by one-way ANOVA followed by Tukey’s multiple comparison test (* *p* < 0.05; ** *p* < 0.01; *** *p* < 0.001).

### AZD1390 protects against neuronal cell death in *ex vivo* brain slice models of stroke

After reproducing the ability of ATM inhibitors to protect against ferroptosis of cancer cells, we wish to extend such a finding to non-cancer settings, such as cortical neuronal cell death induced by ischemic stroke. To assess the pharmacological effectiveness of ATM inhibitors, we employed organotypic brain slice culture models of stroke by exposing them to OGD. Treatment with three different ATM inhibitors to brain slices of Sprague-Dawley (SD) rats following OGD consistently demonstrated that these inhibitors effectively protect against neuronal cell death under ischemic conditions (Fig 2A-D). AZD1390 rescued neuronal cell viability in this assay, although its effectiveness was somewhat lower compared to other ATM inhibitors. However, given that AZD1390 is a clinical compound demonstrating benefits in promoting axon regeneration and functional recovery following spinal cord injury ^16^, we focused subsequent studies on this compound. To verify whether AZD1390 has its protective effect via the ferroptotic pathway, we determined transcription levels of ferroptosis markers, xCT (also known as Solute Carrier Family 7 Member 11 (*SLC7A11*)) and prostaglandin-endoperoxide synthase 2 (*PTGS2*), from rat brain slices after transient OGD treatment. The transcript levels of both xCT and PTGS2 were elevated in brain slices following OGD treatment. However, AZD1390 reduced these levels in a dose-dependent manner, mirroring the effects of the ferroptosis inhibitor, ferrostatin-1 (Fer-1) (Fig 2E). We also evaluated the effectiveness of AD1390 in reducing neuronal death cortical brain slices from C57BL/6J (B6/J) mice and observed that, as with the rat brain slices, AZD1390 treatment led to increased cell viability (Fig 2F). These findings suggest that neuronal cell death in both rat and mouse brain slices is mediated by ferroptosis, and AZD1390 effectively protects against this process.

**Figure 2.**
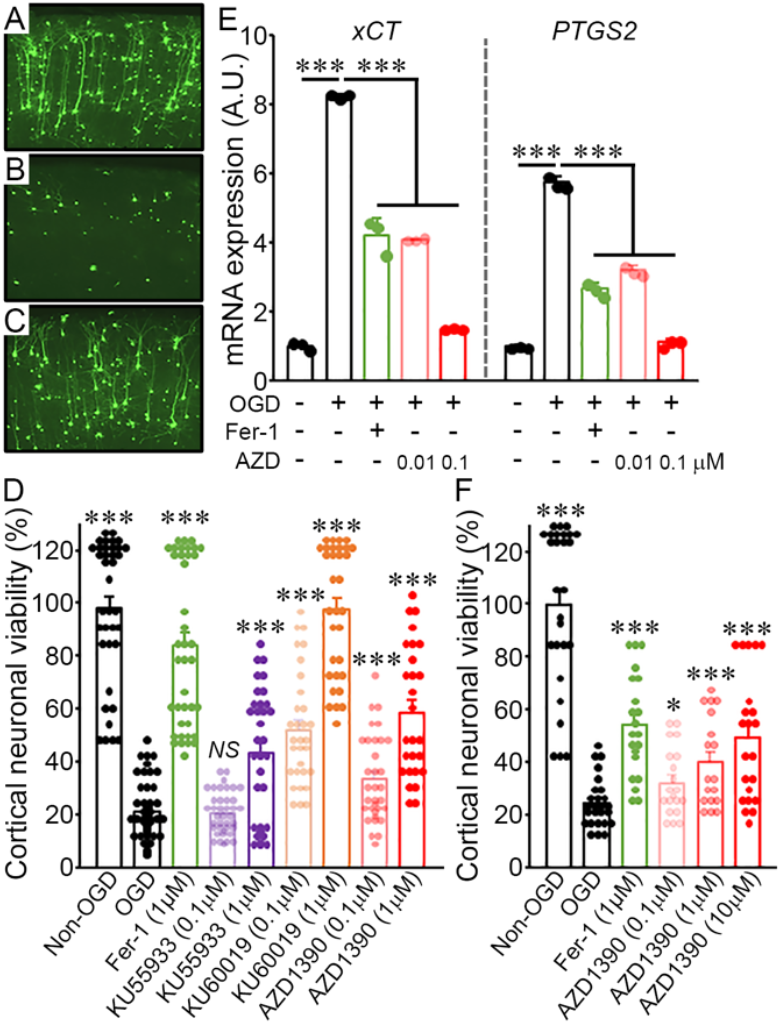
AZD1390 increases neuronal cell viability in the cortical brain slice ischemia assay. **(A – C)** Representative images showing the number of YFP-transfected cortical pyramidal neurons in rat brain slices under different conditions: normoxia (A), after 4.5 minutes of OGD (B), and after 4.5 minutes of OGD followed by treatment with ferrostatin-1 (Fer-1, 1μM) (C). **(D)** The graph indicating the average cell viability. Treatment with the ferroptosis inhibitor Fer-1 and ATM inhibitors, including AZD1390, significantly increased cell viability. **(E)** AZD1390 reduced the induction of ferroptosis markers, xCT and PTGS2, in the rat brain slice model of ischemic stroke. Transcript levels of xCT and PTGS2 were measured using qPCR from the rat brain slices. **(F)** Neuronal cell viability in mouse brain slices increased in a dose-dependent manner with AZD1390 treatment. Data represent the mean ± SEM and statistical significance was determined by one-way ANOVA followed by Tukey’s multiple comparison test (* *p* < 0.05; ** *p* < 0.01; *** *p* < 0.001 *vs*. OGD).

### Brain-penetrant AZD1390 reduces infarct volume after pMCAO in an *in vivo* stroke model

To extend these findings *in vivo*, we induced an ischemic stroke by surgical occlusion of the middle cerebral artery of two inbred strains of mice, B6/J and C57BL/6NJ (B6/NJ), and determined the effects of administered AZD1390. Both mouse strains received 20 mg/kg of AZD1390 via oral gavage two hours after permanent MCAO, and infarct volume was measured 72 hours post-pMCAO (Fig 3A). This dosing strategy follows a prior study demonstrating AZD1390’s brain penetration in primate and mouse brains ^17^, and delivery at two hours after surgical occlusion is consistent with a translationally feasible therapeutic approach. AZD1390, when compared with vehicle control, was found to significantly reduce infarct volume in both mouse strains, B6/J and B6/NJ, with decreases of 55.6% in B6/J and 37.1% in B6/NJ (Fig 3B and C). Next, we investigated the levels of malondialdehyde (MDA), an end-product of lipid peroxides during ferroptosis. Ischemic mice showed dramatically reduced levels of MDA after AZD1390 treatment compared to vehicle-treated mice (Fig 3D and E). Collectively, these data show that in an ischemic model stroke, the reduction of infarct volume by AZD1390 was associated with inhibition of ferroptosis.

**Figure 3.**
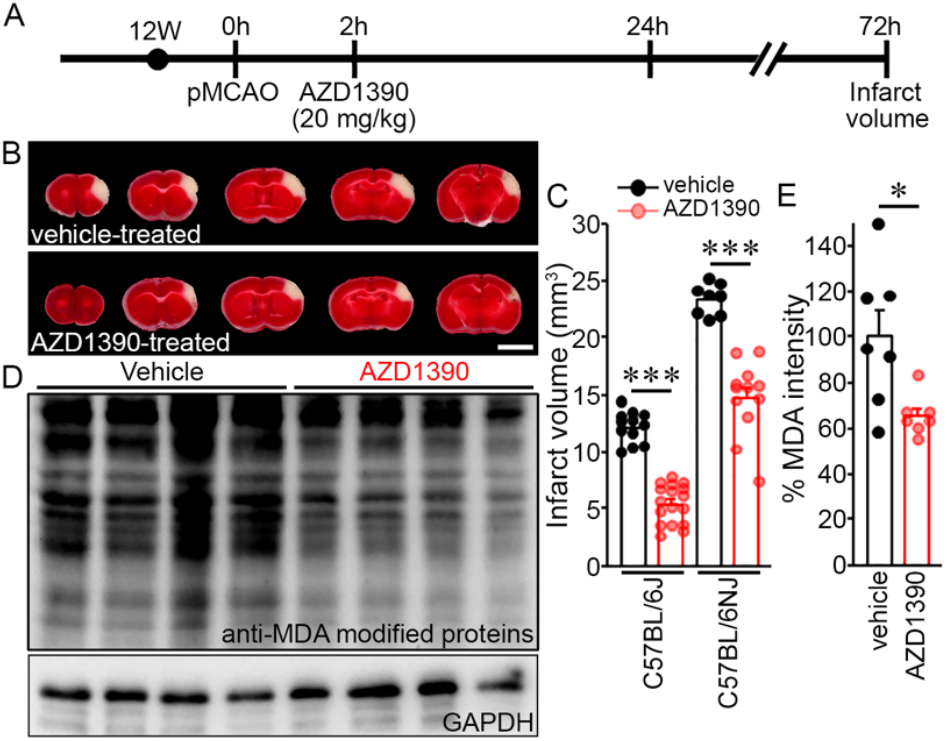
AZD1390 reduces infarct volume after pMCAO in an *in vivo* stroke model. **(A)** Schematic timeline illustrating the experimental procedure, including ischemic stroke induction through pMCAO and AZD1390 treatment. **(B)** Serial brain slice sections (1 mm) from the B6/NJ inbred mouse strain. Infarcted tissue appears as white after 2% TTC staining. Scale bar: 5 mm. **(C)** The graph showing infarct volume 72 hours after pMCAO. AZD1390 treatment significantly reduced infarct volume compared to vehicle-treated group. **(D)** Western blots analysis of malodialdehyde (MDA), an end-product of lipid peroxides during ferroptosis, in brain tissues from mice subjected to pMCAO followed by vehicle or AZD1390 treatment. GAPDH was used as a loading control. **(E)** Quantification of MDA levels normalized to GAPDH. Data represent the mean ± SEM and statistical significance was determined by two-tailed Student’s *t*-test (* *p* < 0.05; *** *p* < 0.001).

## Discussion

This study provides evidence for the role of ferroptosis as a mechanism underlying neuronal cell death in ischemic stroke, and identifies ATM inhibition as a potential therapeutic strategy. While ferroptosis has been previously implicated in various pathological conditions, including cancer and acute kidney injury, our findings highlight its critical role in ischemic stroke-induced neuronal damage ^1,2,5,6^. By demonstrating that ATM inhibitors, particularly the brain-penetrant AZD1390, mitigate neuronal cell death and infarct volume in both *ex vivo* and *in vivo* models, we establish a direct link between ischemic stroke pathology and the ferroptotic pathway. The downregulation of ferroptosis markers, such as xCT and PTGS2, further supports our discovery that ATM inhibition mediates its protective effect by inhibiting ferroptosis. These results extend our understanding of the molecular events contributing to neuronal injury and open a new direction in stroke therapy by targeting ferroptosis. Furthermore, the reduction in infarct volume in mouse models treated with AZD1390 two hours after ischemic stroke occlusion emphasizes its potential as a clinically relevant intervention. The translation potential of AZD1390 is oral administration and the ability to cross the blood-brain barrier ^16,17^, making it a strong candidate for translational application. As AZD1390 is now used in clinical trial (NCT03423628).

This study elevates ATM inhibition as a novel therapeutic target for ischemic stroke and emphasizes the translational potential of targeting ferroptosis to mitigate neuronal damage and improve outcomes for patients suffering from ischemic stroke.

## Conclusions

This study establishes ferroptosis as a key mechanism underlying neuronal cell death in ischemic stroke and highlights ATM inhibition, particularly with the brain-penetrant AZD1390, as a promising therapeutic strategy. By demonstrating that ATM inhibitors alleviate ferroptosis-induced neuronal cell death and reduce infarct volume in both *ex vivo* and *in vivo* models, this research provides compelling evidence for targeting ferroptosis as a novel approach to stroke treatment. The efficacy of AZD1390, an orally bioavailable and brain-penetrating clinical compound, underscores its translation relevance. These findings pave the way for the development of ferroptosis-targeting therapies to mitigate neuronal injury and improve clinical outcomes for stroke patients, offering a pathophysiologically grounded approach with clinical feasibility.

## Acknowledgements

The authors thank AstraZeneca for providing AZD1390 through the AstraZeneca Open Innovation Programme.

## References

1. Dixon SJ, Lemberg KM, Lamprecht MR, Skouta R, Zaitsev EM, Gleason CE, Patel DN, Bauer AJ, Cantley AM, Yang WS, et al. Ferroptosis: an iron-dependent form of nonapoptotic cell death. Cell. 2012. 149(5):1060–72. doi: 10.1016/j.cell.2012.03.042.

2. Yang WS and Stockwell BR. Synthetic lethal screening identifies compounds activating iron-dependent, nonapoptotic cell death in oncogenic-RAS-harboring cancer cells. Chemistry & Biology. 2008. 15(3):234–45. doi: 10.1016/j.chembiol.2008.02.010.

3. Yang WS, SriRamaratnam R, Welsch ME, Shimada K, Skouta R, Viswanathan VS, Cheah JH, Clemons PH, Shamji AF, Clish CB, et al. Regulation of ferroptotic cancer cell death by GPX4. Cell. 2014. 156(1-2):317-31. doi: 10.1016/j.cell.2013.12.010.

4. Doll S, Freitas FP, Shah R, Aldrovandi M, Costa da Silva M, Ingold I, Grocin AG, Xavier da Silva TN, Panzilius E, Scheel CH, et al. FSP1 is a glutathione-independent ferroptosis suppressor. Nature. 2019. 575(7784):693–698. doi: 10.1038/s41586-019-1707-0.

5. Bu ZQ, Yu HY, Wang J, He X, Cui YR, Feng JC, Feng J. Emerging Role of Ferroptosis in the Pathogenesis of Ischemic Stroke: A New Therapeutic Target? ASN Neuro. 2021. 13:17590914211037505. doi: 10.1177/17590914211037505.

6. Alim I, Caulfield JT, Chen Y, Swarup V, Geschwind DH, Ivanova E, Seravalli J, Ai Y, Sansing LH, Ste Marie EJ, Hondal RJ, et al. Selenium Drives a Transcriptional Adaptive Program to Block Ferroptosis and Treat Stroke. Cell. 2019. 177(5):1262-79.e25. doi: 10.1016/j.cell.2019.03.032.

7. Chen PH, Wu J, Ding CC, Lin CC, Pan S, Bossa N, Xu Y, Yang WH, Mathey-Prevot B, Chi JT. Kinome screen of ferroptosis reveals a novel role of ATM in regulating iron metabolism. Cell Death Differ. 2020. 27(3):1008–22. doi: 10.1038/s41418-019-0393-7.

8. Wu H, Liu Q, Shan X, Gao W, Chen Q. ATM orchestrates ferritinophagy and ferroptosis by phosphorylating NCOA4. Autophagy. 2023. 19(7):2062–2077. doi: 10.1080/15548627.2023.2170960.

9. Chen PH, Tseng WHS, Chi JT. The Intersection of DNA Damage Response and Ferroptosis-A Rationale for Combination Therapeutics. Biology (Basel). 2020. 9(8):187. doi: 10.3390/biology9080187.

10. Jucaite A, Stenkrona P, Cselényi Z, De Vita S, Buil-Bruna N, Varnäs K, Savage A, Varrone A, Johnström P, Schou M, et al. Brain exposure of the ATM inhibitor AZD1390 in humans—a positron emission tomography study. Neuro Oncol. 2020. 23(4):687–696. doi: 10.1093/neuonc/noaa238.

11. Lee HK, Keum S, Sheng H, Warner DS, Lo DC, Marchuk DA. Natural allelic variation of the IL-21 receptor modulates ischemic stroke infarct volume. J Clin Invest. 2016. 126, 2827–2838. doi: 10.1172/JCI84491.

12. Lee HK, Koh S, Lo DC, Marchuk DA. Neuronal IL-4Rα modulates neuronal apoptosis and cell viability during the acute phases of cerebral ischemia. FEBS J. 2018. 285, 2785–2798. doi: 10.1111/febs.14498.

13. Lee HK, Widmayer SJ, Huang MN, Aylor DL, Marchuk DA. Novel Neuroprotective Loci Modulating Ischemic Stroke Volume in Wild-Derived Inbred Mouse Strains. Genetics. 2019. 213(3):1079–1092. doi: 10.1534/genetics.119.302555.

14. Lee HK, Kwon DH, Aylor DL, Marchuk DA. A cross-species approach using an in vivo evaluation platform in mice demonstrates that sequence variation in human RABEP2 modulates ischemic stroke outcomes. Am J Hum Genet. 2022. 109(10):1814–1827. DOI: 10.1016/j.ajhg.2022.09.003.

15. Talele S, Zhang W, Chen J, Gupta SK, Burgenske DM, Sarkaria JN, Elmquist WF. Central Nervous System Distribution of the Ataxia-Telangiectasia Mutated Kinase Inhibitor AZD1390: Implications for the Treatment of Brain Tumors. J Pharmacol Exp Ther. 2022 383:91–102. doi: 10.1124/jpet.122.001230.

16. Ahmed Z and Tuxworth RI. The brain-penetrant ATM inhibitor, AZD1390, promotes axon regeneration and functional recovery in preclinical models of spinal cord injury. Clin Transl Med. 2022. 12:e962. doi: 10.1002/ctm2.962.

17. Durant ST, Zheng L, Wang Y, Chen K, Zhang L, Zhang T, Yang Z, Riches L, Trinidad AG, Fok JHL, et al. The brain-penetrant clinical ATM inhibitor AZD1390 radiosensitizes and improves survival of preclinical brain tumor models. Sci Adv. 2018. 4(6):eaat1719. doi: 10.1126/sciadv.aat1719.

