## Supplemental materials and methods for "Targeting the ferroptosis pathway: A novel compound, AZD1390, protects the brain after ischemic stroke"

**Animals.** All inbred mouse strains were obtained from the Jackson Laboratory (Bar Harbor, ME), and then bred locally to generate mice used in all experiments. Sprague-Dawley (SD) rats for *ex vivo* brain slice culture experiments were obtained from Charles River (Wilmington, MA). 12 ± 1 week-old mice (both male and female animals) were used for pMCAO and postnatal day 7 mice and rats were used for *ex vivo* brain slice culture. All animal study procedures were conducted under protocols approved by the Duke University IACUC in accordance with NIH guidelines.

**Cell culture.** MDA-MB-231 cells were obtained from the Cell Culture Facility at Duke University. Before freezing, the cell lines were authenticated via STR DNA profiling and confirmed to be free of mycoplasma contamination. The cells were maintained for less than six months and cultured in DMEM medium (GIBCO-11995) supplemented with 10% heat-inactivated fetal bovine serum (ThermoFisher #10082147) and antibiotics (streptomycin (10,000 U/ml) and penicillin (10,000 U/ml); ThermoFisher #15140122). Cultures were incubated at 37°C in a humidified atmosphere with 5% $CO_2$.

**AZD1390 preparation.** Pharmaceutical-grade AZD1390 for *in vivo* studies was obtained directly from AstraZeneca and was prepared as recommended by the manufacture's formulation and doses for *in vivo* mouse studies. Briefly, a dose of 20 mg/kg was suspended in a vehicle consisting of 0.5% hydroxypropyl methylcellulose (HPMC) and 0.1% Tween 80 in water. This suspension was either stirred or rotated overnight at room temperature in the absence of light. Freshly prepared drug formulations were used within 2 days for *in vivo* experiments.

**Chemicals.** All chemicals, including drugs and inhibitors, were used as previously described [1]. Briefly, erastin (Bio-Techne #5499), ferrostatin-1 (Cayman #17729), deferoxamine (Sigma D9533), liproxstatin-1 (Cayman #17730), KU55933 (Selleckchem #S1092), and KU60019 (Selleckchem #S1570) were prepared as stock solutions in DMSO. For experiments, these stock solutions were diluted 1:1000 in culture media to prepare for working solutions.

**Cell Viability and Cytotoxicity Assays.** Cell viability was assessed using the CellTiter-Glo luminescent viability assay (Promega) as recommended by the manufacturer's instructions. In brief, 15 µl of the CellTiter-Glo substrate was added to wells containing cells in 100 µl of culture medium in a 96-well plate. The plate was shaken for 10 minutes, and the luminescent signal was quantified using a chemiluminescence plate reader. Cytotoxicity was evaluated by CellTox Green assay (Promega), which was diluted by the ratio of 1:1000 to the culture medium and measuring fluorescence.

**Quantitative PCR.** mRNA levels were measured by quantitative PCR (qPCR) as previously described [2].To quantitative mRNA levels of *xCT* and *PTGS2*, rat brain slices were treated with or without OGD, followed by either Ferrostatin-1 (Fer-1) or AZD1390. All samples were analyzed in triplicate, and an additional assay targeting endogenous *Gapdh* was included to normalize for input cDNA template quantity. Primer sequences for rat included:

*xCT*: Forward 5'- TAC CTG CAG GGC AAT GTG AG-3', Reverse 5'- AGT ATG CCC TTG GGG GAG AT-3'; *PTGS2*: Forward 5'-TGA CTG TAC CCG GAC TGG AT-3', Reverse 5'-CAT GGG AGT TGG GCA GTC AT -3'; *Gapdh*: Forward 5'-GCG AGA TCC CGC TAA CAT CA-3', Reverse 5'- CTT GCG GTC CCC ACA TAC C-3'

**Lipid Peroxidation Assay.** Lipid peroxidation was evaluated using the C11-BODIPY dye (ThermoFisher, D3861). Cells were treated with either vehicle or erastin with/without various dose of AZD1390 for 16 hours. Subsequently, a 10 μM solution of C11-BODIPY was added to the culture medium for 1 hour at 37°C. The cells were harvested, washed, and resuspended in PBS with 1% BSA. Lipid peroxidation levels were analyzed using flow cytometry (FACSCanto II, BD Biosciences).

**Western Blot Analysis.** Protein samples were collected from mouse brain tissues induced ischemic stroke with either vehicle or AZD1390 treatment. Briefly, the protein concentrations were determined via the BCA assay (ThermoFisher, #23227) after lysing pulverized mouse brain samples in RIPA buffers. Approximately 20 µg of protein per sample was resolved on 8% SDS-PAGE gels, transferred to PVDF membranes, blocked with 5% non-fat milk in 1xTBST, and incubated overnight at 4°C with primary antibodies. Antibodies used included MDA (1:1000, ab27642, Abcaam); GAPDH (1:2000, sc-25778, Santa Cruz).

**Preparation of brain slices and oxygen glucose deprivation (OGD).** Preparation of cortical brain slice explants was performed as previously described. [3,4] Coronal brain slices were prepared from P7 rats and mice. Under sterile conditions, brains were dissected and cut into 250 μM coronal slices on a vibratome in chilled culture medium containing 15% heat-inactivated horse serum, 10 mMKCL, 10 mM HEPES, 100U/mL penicillin/streptomycin, 1 mM MEM sodium pyruvate, and 1mM I-glutamaine in Neurobasal A supplemented with 1 μM MK-801.

Brains were divided into hemi-coronal slices. For OGD, slices were suspended at 34°C for 4.5 minutes glucose-free $N_2$-bubbled artificial CSF containing 124 mM NaCl, 25.7 mM

$NaCO_3$, 4.9 mM KCL, 1.2 mM $KPO_4$, 3.1 mM $CaCl_2$, 1.3 mM $MgSO_4$ (pH 7.0). Brain slices were plated into 12-well plates in interface configuration atop solid culture medium made by the addition of 0.5% agarose. After explanting the brain slices, plates were placed for recovery at 30°C for 30 minutes in a humidified incubator under 5% $CO_2$. Gold particle (1.6 μM) coated with plasmids expressing YFP in G-Wiz vector were introduced into the brain slices by biolistic transfection using a Helios Gene Gun (Bio-Rad). Slice cultures were maintained at 30°C for 24 hours in humidified incubator under 5% $CO_2$.

Viable neurons were scored by YFP morphology under fluorescent stereoscope visualization.

**Permanent middle cerebral artery occlusion (pMCAO) and AZD1390 treatment.** Focal cerebral ischemia was induced by direct permanent occlusion of the distal MCA as previously described. [5-7] Briefly, adult mice were anesthetized with ketamine (100 mg/kg) and xylazine (5 mg/kg), and then 0.5% bupivacaine (5 mg/mL) was also administrated by injection at the incision site. The right MCA was exposed by a 0.5 cm vertical skin incision midway between the right eye and ear under a dissecting microscope. After the temporalis muscle was split, a 2 mm burr hole was made with a high-speed micro drill at the junction of the zygomatic arch and the squamous born through the outer surface of the semi-translucent skull. The MCA was clearly visible at the level of the inferior cerebral vein. The inner layer of the skull was removed with fine forceps, and the dura was opened with 32-gauge needle. While visualizing under an operating microscope, the right MCA was electrocauterized. The cauterized MCA segment was then transected with microscissors to verify permanent occlusion. The surgical site was closed with 6-0 sterile nylon sutures, and 0.5 % bupivacaine was applied. The temperature of each mouse was maintained at 37°C with a heating pad during the surgery and then mouse was placed in a recovery chamber (set temperature 37°C) until the animal was fully recovered from the anesthetic. Two hours after pMCAO, animals received a 20 mg/kg dose of AZD1390 via oral gavage using a flexible feeding tube (Instech, Plymouth Meeting, PA). Mice were then returned to their cages and allowed free access to food and water in an air-ventilated room maintained at an ambient temperature of 25°C.

**Infarct volume measurement.** Cerebral infarct volumes were measured 72 hours after distal permanent MCA occlusion as previously described. [5-7] Seventy-two hours after pMCAO surgery, the animals were euthanized, and the brains were carefully removed. The brains were placed in a brain matrix, chilled at -80°C for 4 min to slightly harden the tissue, and then sliced into 1 mm coronal sections. Each brain slice was placed in 1 well of a 24-well plate and incubated for 20 min in a solution of 2% 2,3,5-triphenyltetrazolium chloride (TTC) in PBS at 37°C in the dark. The sections were then washed once with PBS and fixed with 10% PBS-buffered formalin at 4°C. Then 24 hours after fixation, the caudal face of each section was scanned using a flatbed color scanner. The scanned images were used to determine infarct volume. Image-Pro software (Media Cybernetics, Inc., MD) was used to calculate the infarcted area of the hemisphere to minimize error introduced by edema. The total infarct volume was calculated by summing the individual slices from each animal.

**Statistical analyses.** Statistical analyses were performed with GraphPad Prism (GraphPad Software, La Jolla, CA). Significant differences between datasets were evaluated using a two-tailed Student's *t*-test comparisons between two groups or oneway ANOVA followed by Tukey's multiple comparison test for more than two groups. Data are represented as the mean ± SEM. $p < 0.05$ was considered statistically significant. Detailed statistical analyses for all figures are provided in Table S1.

### Supplemental References

1. Lin CC, Yang WH, Lin YT, Tang X, Chen PH, Ding CKC, Qu DC, Alvarez JV, Chi JT. DDR2 upregulation confers ferroptosis susceptibility of recurrent breast tumors through the Hippo pathway. Oncogene. 2021. 40(11):2018–34. doi: 10.1038/s41388-021-01676-x.
2. Lin CC, Lin YT, Chen SY, Setayeshpour Y, Chen Y, Dunn DE, Soderblom EJ, Zhang GF, Filonenko V, Jeong SY, et al. Protein CoAlation on TXNRD2 regulates mitochondrial thioredoxin system to protect against ferroptosis. bioRxiv. 2024. 10.1101/2024.05.16.594391.
3. Lee HK, Keum S, Sheng H, Warner DS, Lo DC, Marchuk DA. Natural allelic variation of the IL-21 receptor modulates ischemic stroke infarct volume. J Clin Invest. 2016. 126, 2827-2838. doi: 10.1172/JCI84491.
4. Lee HK, Koh S, Lo DC, Marchuk DA. Neuronal IL-4Rα modulates neuronal apoptosis and cell viability during the acute phases of cerebral ischemia. FEBS J. 2018. 285, 2785-2798. doi: 10.1111/febs.14498.
5. Lee HK, Widmayer SJ, Huang MN, Aylor DL, Marchuk DA. Novel Neuroprotective Loci Modulating Ischemic Stroke Volume in Wild-Derived Inbred Mouse Strains. Genetics. 2019. 213(3):1079-1092. doi: 10.1534/genetics.119.302555.
6. Lee HK, Kwon DH, Aylor DL, Marchuk DA. A cross-species approach using an in vivo evaluation platform in mice demonstrates that sequence variation in human RABEP2 modulates ischemic stroke outcomes. Am J Hum Genet. 2022. 109(10):1814-1827. DOI: 10.1016/j.ajhg.2022.09.003.

7. Lee HK, Wetzel-Strong SE, Aylor DL, Marchuk DA. A neuroprotective locus modulates ischemic stroke infarction independent of collateral vessel anatomy. Front Neurosci. 2021. 15:705160. DOI: 10.3389/fnins.2021.705160.
